# Molecular docking-based screening for novel inhibitors of the human immunodeficiency virus type 1 protease that effectively reduce the viral replication in human cells

**DOI:** 10.1101/2020.11.14.382895

**Authors:** Carla Mavian, Roxana M Coman, Xinrui Zhang, Steve Pomeroy, David A. Ostrov, Ben M Dunn, John W. Sleasman, Maureen M Goodenow

## Abstract

Therapeutic pressure by protease inhibitors (PIs) contributes to accumulation of mutations in the HIV type 1 (HIV-1) protease (PR) leading to development of drug resistance with subsequent therapy failure. Current PIs target the active site of PR in a competitive manner. Identification of molecules that exploit non-active site mechanisms of inhibition is essential to overcome resistance to current PIs. Potential non-active site HIV-1 protease (PR) inhibitors (PI) were identified by in silico screening of almost 140,000 molecules targeting the hinge region of PR. Inhibitory activity of best docking compounds was tested in an in vitro PR inhibition biochemical assay. Five compounds inhibited PR from multiple HIV-1 subtypes in vitro and reduced replicative capacity by PI-sensitive or multi-PI resistant HIV-1 variants in human cells ex vivo. Antiviral activity was boosted when combined with Ritonavir, potentially diminishing development of drug resistance, while providing effective treatment for drug resistant HIV-1 variants.

## Introduction

With no available cure, treatment of HIV infection requires life-long therapy. The landscape of HIV treatment has significantly benefited from the approval of multidrug combinations of antiretroviral therapies (cART) based on several new antiretroviral drugs (ARVs) with enhanced potency, safety and tolerability. Since introduction of ARVs for pre-exposure prophylaxis (PrEP) in 2012 as a measure of HIV prevention, HIV transmission has substantially decreased (1, 2). However, every year approximately 2 million new HIV infections are reported worldwide, of which 50,000 are in the U.S. (3). Effective strategies to prevent HIV transmission are needed as successful implementation and worldwide breadth of PrEP represent still a challenge. HIV research attention remains focused on the developing long-acting ARVs to mitigate drug adherence and resistance issues.

Among the pharmacologic agents undergoing evaluation for use as PrEP, earlier phase studies are currently evaluating protease inhibitors (PIs) for intravaginal rings (4). Inhibiting PR prevents maturation of viral progeny, compromises further entry and inhibits viral spread (5, 6). Including PIs in cART has significantly improved length and quality of life of HIV infected individuals (5, 7). However, currently PIs in use present side effects of antiretroviral drugs, development of drug resistance, and transmission of multidrug resistant strains in 9% to 15% of new infections represent major challenges for successful cART (8–10). HIV type 1 (HIV-1) PR is a homodimeric enzyme of 99 amino acids [aa] essential for HIV replication (6). Although, inhibition of HIV-1 PR enzymatic activity by competitive inhibition of substrate binding in the active site has pleiotropic effects on the viral life cycle (6), one of the major drawbacks is the emergence of viral species exhibiting resistance to PIs that limits long-term treatment options. Design of PIs efficient against multi-drug resistant PR is fundamental to overcome development of resistance. Novel mechanisms of inhibiting PR target the flexibility of the enzyme in non-active sites, such as the elbow/hinge region of the flaps (aa 39-42), the tip of the flap (aa 43-58), or the β-sheet region (aa 1-5, 95-99) (11–13). The flaps close to initiate processing once substrate binds in the active site of PR. Inhibition of the flaps would theoretically change the conformation of PR and modulate the active site (14). The nature of the interaction of non-active site PIs with the PR may also decrease the therapeutic index of current PIs by reducing dosage or frequency of administration, and potentially minimizing side effects. Previously, nonactive site PIs have been identified and tested against HIV-1 PR in biochemical assays, although evidence for effectiveness reduction of HIV-1 infection in human cells is lacking (11, 13, 15). We explored the hypothesis that compounds selected by *in silico* binding to the hinge region of PR are effective as inhibitors of viral replication, including multi-PI resistant viruses, in human cells.

## Materials and methods

### Molecular Docking

The whole repository of 139,739 small molecules (molecular weight, <500) of the National Cancer Institute/Developmental Therapeutics Program (NCI/DTP) was screened for identification of novel PIs (16). The three-dimensional coordinates of the 139,739 NCI/DTP compounds were obtained in MDL SDF-format and converted to mol2 format with SDF2MOL2 (UCSF). Molecular docking was performed with DOCK v5.1.0 (i) selecting for structural pockets in HIV-1 PR suitable for interactions with drug-like small molecules and (ii) performing molecular docking simulations where each small molecules is positioned in the selected structural pocket and scored based on predicted polar (H-bond) and nonpolar (van der Waals) interactions (17). Docking calculations were performed with the 15 October 2002 development version of DOCK v5.1.0 (18).

The crystal structure of HIV protease (PDB code 1TW7) was used the basis for molecular docking (19). Molecular docking was realized on conserved residues located on the side of the PR grove formed by the elbow of the flap and the 60’s loop (43-58 aa) (**Fig S1**). DOCK explores the molecular surface using sets of spheres as a guide to search for orientations of each molecule that fit into the selected sites to describe potential binding pockets. The sites selected for molecular docking were defined using the SPHGEN program and filtered through the CLUSTER program (20). Seventy-two spheres were used to define the HIV-1 PR site for molecular docking. Each compound in the NCI/DTP database was positioned in the selected site in 100 different orientations at the University of Florida High Performance Computing Center. Intermolecular AMBER energy scoring (van der Waals_columbic), contact scoring, and bump filtering were implemented in DOCK v5.1.0. PYMOL (https://www.pymol.org/) was used to generate molecular graphic images (21).

### In vitro PR inhibition assays

The compounds were obtained from the NCI/DTP and solubilized in DMSO (Sigma) for use in HIV-1 PR inhibition assays. Cloning, expression, and purification procedures of HIV-1 PRs obtained from patent samples has been previously described elsewhere (22–24). For detailed information of mutations harbored in the multi PI-resistant HIV-1PR as compared to the PI-sensitive PR, full amino acid sequences is given in supplementary materials (**Fig S2**). HIV-1 PRs subtype variants were assayed kinetically for inhibitory activity at 80 μM at 37 °C using selected compounds as previously described (22).

### Cell culture

TZM-bl cells (ARP5011) with an integrated luciferase gene under the control of the HIV-1 promoter were obtained through the NIH AIDS Research and Reference Reagent Program, Division of AIDS (25, 26). TZM-bl cells were cultured in DMEM media (Gibco) containing 10% fetal human serum (FHS), 2 mM L-Glutamine, 0.05% sodium bicarbonate and 0.1 mg/ml Penicillin and Streptomycin. Peripheral blood mononucleated cell (PBMCs) were obtained from Lifesouth Community Blood Center, Gainesville, FL. with approval for use of human cells by the Institutional Review Board. PBMCs were stimulated with phytohemaglutinin (2mg/ml) for 72 hours and cultured in RPMI Medium (Gibco) with 10% FHS. Both cell types were kept in an incubator at 37°C with 5% CO_2_.

### HIV-1 PI-sensitive and multi PI-resistant variants

Virus stocks of the CCR5 using, replication competent PI-sensitive (HIV-1_AD_) or multi (Ritonavir (RTV) and Indinavir) PI-resistant (HIV-1 AD02) HIV-1 variants were prepared by transfecting plasmid DNA from a molecular clone into 293T cells and titered as described previously (27). Briefly, the molecular clone pAD8 (Theodore et al, 1996), a stable full-length macrophage-tropic HIV type 1 molecular clone, was used to generate a protease inhibitor-resistant molecular clone: a viral isolate from a PR drug-resistant patient was used to generate amplified DNA gag-pol amplicons as previously described by our laboratory (28, 29). The drug-resistant PR was obtained from the replication competent virus present in the peripheral blood of a patient who had failed PIs (28, 29). The gag-pol regions carried by the PI-sensitive (HIV-1_AD_) and the multi PI-resistant (HIV-1_AD02_) HIV-1 variants have been well characterized by our laboratory (28) (**Fig S2**).

### Cell viability assays

Viability in presence of compounds was measured at day 4 or 7, TZM-bl cells or PBMCs, using CellTiter 96 AQueous one solution cell proliferation assay (Promega) following manufacturer instructions. Toxicity of compounds was defined as a concentration that produced more than 10% decline in viability of TZM-bl cells or PBMCs were viable at days 4 or 7 of exposure, respectively, compared with DMSO-treated controls (28, 30, 31).

### HIV-infection and anti-viral activity assays

Anti-viral activity was evaluated *ex vivo* by adding the highest non-toxic concentration of compound or 25 uM RTV (Sigma) solubilized in DMSO on the day of infection by PI-sensitive (HIV-1_AD_) or multi (RTV and IDV) PI-resistant (HIV-1_AD02_) HIV-1 variants at 25 or 45 TCID50 per ml for PBMCs or TZM-bl cells, respectively. Replication was determined on post-infection day 7 in PBMC by measuring supernatant p24 antigen levels (Perkin-Elmer), or day 4 in TZM-bl cells by measuring luciferase activity using the luciferase assay system (Promega, Madison, WI) and the microplate luminometer Monolight 3096 (BD Biosciences, San Jose, CA). Inhibitory concentration (IC)50 was calculated based on anti-viral activity in TZM-bl cells with a nonlinear regression analysis (GraphPad Prism 6 Software). Reduction of viral replication by more than 50% was considered successful anti-viral activity. Combinatorial effect of drug with RTV was tested at IC_50_, and higher or lower concentrations than IC_50_.

### Statistical analysis

Linear mixed models with Bonferroni correction to adjust for multiple comparisons were performed with SAS/STAT 9.4 Software.

## Results

### Identification of compounds that inhibit the PR through targeting PR non-active sites

We explored the hypothesis that compounds selected *in silico* for binding to the hinge region of PR are effective as inhibitors of HIV-1 replication, including multi-PI resistant viruses, in human cells. Our of 139,739 compounds, the top 40 highest-scoring compounds targeting the structural pockets in the hinge of the PR (**Fig S2**) were obtained from the NCI/DTP for use in HIV PR inhibition assays. Compounds were initially tested *in vitro* at 80 μM to screen for inhibitory activity against PR subtypes A, B or C. Compounds inhibiting at least one subtype of the PR were considered as candidate compounds for further experiments. Out of the 40 top scoring compounds, 12 compounds inhibited subtype B PR; 11 also inhibited subtype C PR, while 8 inhibited subtype A PR. Compounds that have been identified as PIs candidates are reported in **Table 1**.

**Table 1.**
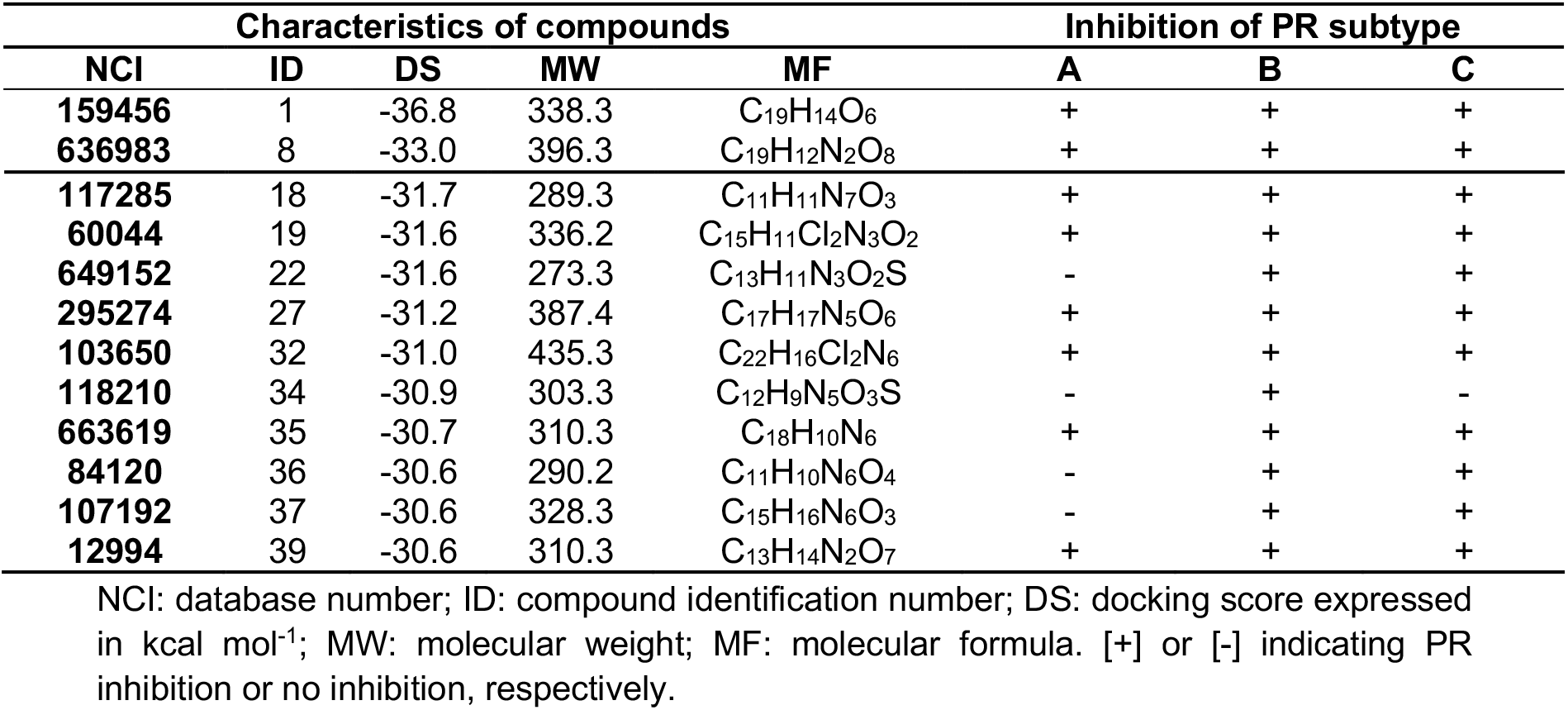
Chemical characteristics and inhibitory activity of selected compounds.

Afterwards, we tested the 12 compounds for toxicity in human cells. A compound was considered toxic when that produced more than 10% decline in viability of TZM-bl cells or PBMCs at a certain concentration (**Table S1**). In TZM-bl cells, compound 19 was toxic at 200 μM reducing cell viability of 28% at 200 μM, compounds 32 and 35 were toxic at concentrations higher than 25 μM reducing cell viability of more than 50%, while nine compounds [compounds 1, 8, 18, 22, 27, 24, 36, 37 and 39] were non-toxic at 200 μM. Compounds showed greater toxic effect in PBMCs: while only four compounds [compounds 8, 34, 36, 37 and 39] were non-toxic at 200 μM; compounds 18 or 22 were toxic at 200 μM reducing cell viability of 28% or 20%, respectively; compounds 19, 32 or 35 were toxic at 50 μM, reducing cell viability of 24%, 32% or 29%, respectively; while compound 27 reduced cell viability of 50% at 25 μM. Compound 1 did not show toxicity in TZM-bl cells, however reduced viability of PBMC by 28% at 10 μM and by 30% at higher concentrations, and therefore was not considered in further testing.

### Anti-viral activity of five lead compounds as novel PIs

Screening for anti-viral activity was evaluated *ex vivo* by adding the compound at the highest non-toxic concentration on the day of infection with either PI-sensitive or multi-PI-resistant HIV-1 variants (**Fig 1**). In PBMC, compounds 27 and 36 reduced replication of PI-sensitive HIV-1_AD_ by more than 60%, while compounds 18, 22, or 35 reduced replication 80% to 90% (**Fig 1A**). Replication of multi-drug resistant HIV-1_AD02_ was reduced by several compounds to levels that were greater than PR sensitive virus, but similar to levels of inhibition by RTV, classic active site PI (**Fig 1A**). Although compounds were less effective against PR resistant virus in TZM-bl cells, compounds 18, 27 or 32 reduced replication by both viruses more than 80% (**Fig 1B**). Some compounds showed discordant performance in PBMCs and TZM-bl cells, and therefore in our further investigation, we analyzed compounds that presented the same trend in both cells: five compounds (18, 22, 27, 32 and 35) inhibited HIV-1 variants in human primary lymphocytes and TZM-bl cells. The IC_50_ of 18 or 27 for either HIV-1 variant was almost identical, being 62.7 μM and 68.5 μM [compound 18] and 26.0 μM and 27.1 μM [compound 27], for HIV-1_AD_ or HIV-1_AD02_ respectively. IC_50_ of compounds 22, 32 and 35 were 53.5 μM or 71.9 μM, 3.8 μM or 13.7 μM, and 1.4 μM or 6.3 μM for HIV-1_AD_ or HIV-1_AD02_, respectively; that is 1.3-fold, 3.6-fold or 4.5-fold lower for HIV-1_AD_ as compared to HIV-1_AD02_, respectively. As previous allosteric PIs (11–13), the five lead compounds that we report presented aromatic structures (**Fig S3**).

**Figure 1.**
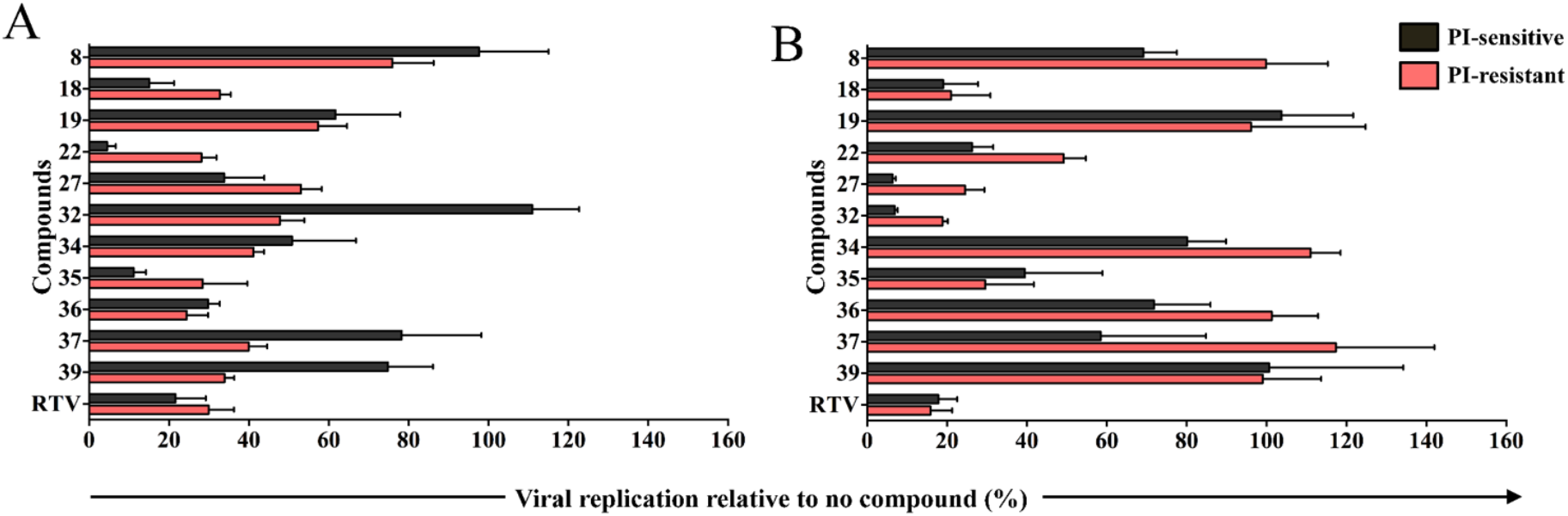
Anti-viral activity screening of compounds in human cells. (A) PBMCs or (B) TZM-bl cells infected with HIV-1_AD_ (black) or HIV-1_AD02_ (red) in presence of compounds. Data are expressed as mean and standard deviation.

Combinatorial effect of drug was tested combining 25 μM RTV and higher or lower concentrations than IC_50_ of lead compounds. Combinations of compounds 18, 22, 27, 32 or 35 with 25 μM RTV were non-toxic in TZM-bl cells. Combination with RTV increased antiviral activity of all compounds against both PI-sensitive and PI-resistant HIV-1 variants (**Fig 2**). Combinations of RTV with lead compounds were more effective than 25 μM RTV alone against either HIV-1_AD_ or HIV-1_AD02_: RTV with compound 32 at 10 μM, compound 35 at 25 μM, or compounds 22 or 27 at 50 and 100 μM respectively, were more effective than 25 μM RTV alone (**Fig 2; Table S2**). While no compound alone inhibited HIV-1 variants to the extent of inhibition by 25 μM RTV, combinations of RTV with compound 18 at 50 and 100 μM were more effective than 25 μM RTV alone against HIV-1_AD02_ (**Fig S1; Table S2**).

**Figure 2.**
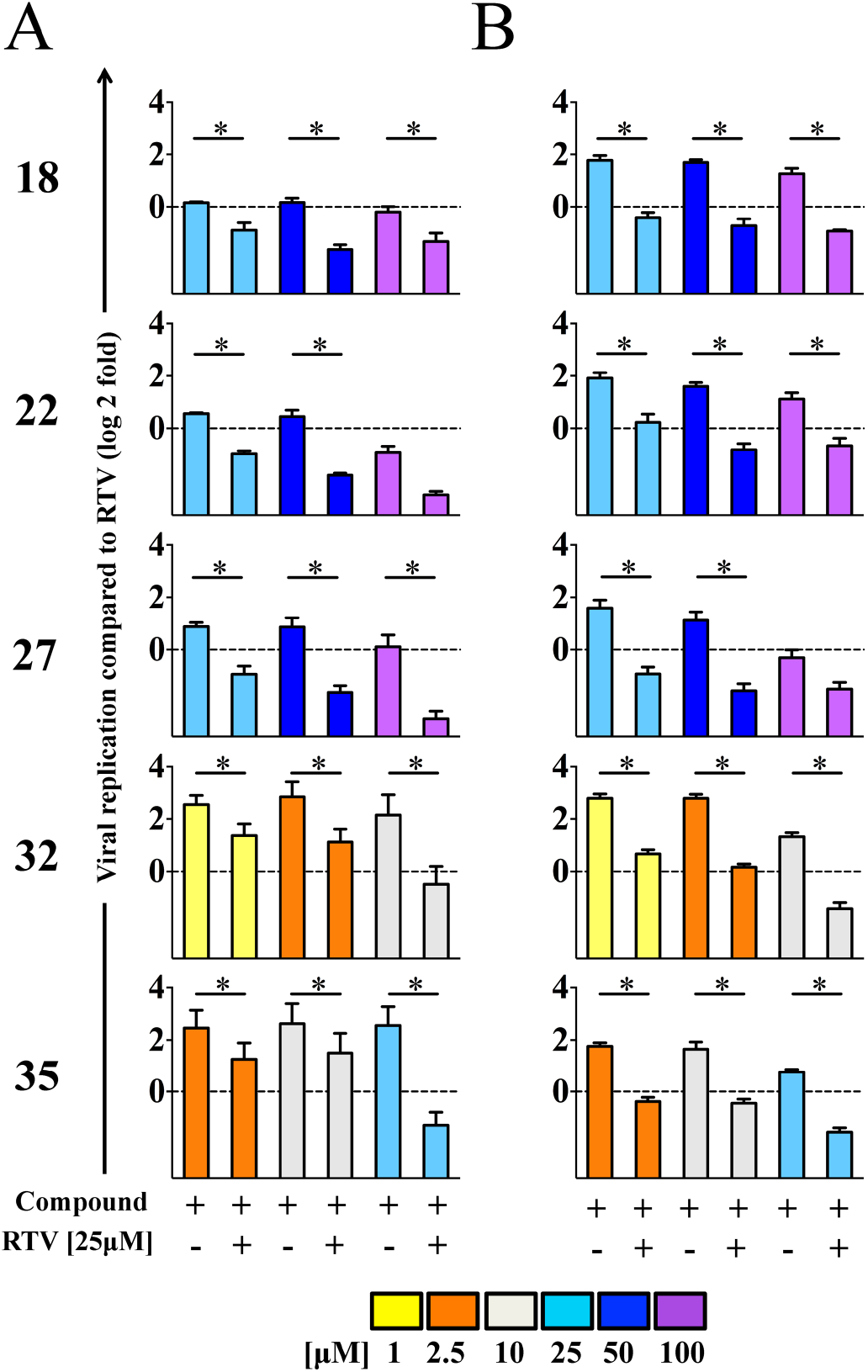
Anti-viral activity of lead compounds in combination with RTV. Anti-viral activity against HIV-1_AD_ (A) or HIV-1_AD02_ (B) of compounds alone or in combination with 25 μM RTV expressed as fold-change relative to RTV (dotted line). Data are expressed as mean and standard deviation. Linear mixed models with Bonferroni correction were performed to adjust for multiple comparisons (SAS/STAT 9.4). Asterisks show statistically significant differences (p<0.05/9=0.0055).

## Discussion

Long-term success of HIV therapy requires both efficacy and safety/tolerability and novel drug development for patients with drug resistance. HIV-1 resistance to anti-retroviral drugs develops with suboptimal levels of inhibitor and is associated with high risk of therapy failure (6). Discovery of drugs that exploit alternative mechanisms of HIV inhibition is essential to avoid or prevent further development of drug-resistance. Our study presents a set of molecules that have similar antiviral activity against PI-resistant and PI-sensitive HIV-1 variants, both *in vitro* and *ex vivo* assays, consistent with a non-active site targeting mechanism. These non-active site PIs were able to inhibit HIV-1 infection by molecular clones generated *ex vivo* from PI-sensitive and PI-resistant naturally occurring HIV-1 variants.

This is the first report of novel PIs that are effectively reducing HIV-1 infection in human cells. PIs that target sites other than the active site in PR would overcome current drug-resistant variants by not competing with natural substrates, and their effect would be undiminished by higher concentrations of substrate (12, 15). Our study design based on *in silico* modeling to find novel PIs that target the non-active site of HIV-1 PR, is supported by *in vitro* studies with similar compounds defined as allosteric PIs (32). Recently, a hinge PI was reported to act by a non-competitive mechanism when delivered together with an active site drugs (13). However, our study presents two caveats. The compounds we describe are a first generation compound that requires further optimization. Further structure-activity modulation of these compounds would lead to the development of more effective molecules with an anti-viral activity at concentrations that would be suitable for pharmaceutical use. Second, preliminary attempts to obtain a crystal structure of the HIV-1 protease bound to the compounds failed. This suggests that the binding of the compounds destabilizes the enzyme inducing a conformation, which differs from the classical PI binding, as reported as well by Tiefenbrunn *et al*, which would be in agreement with our simulation (11). Despite the PR-compounds crystals are not available, our results are consistent with non-active site mechanism of inhibition. The similar anti-viral activity against PI-sensitive and PR-resistant HIV-1 variants suggests that the PR of the PI-resistant variant is naïve to the inhibitory mechanism exploited by the compounds. Indeed, the RTV boosting effect we observed when combining RTV with the lead compounds, is also consistent with, but not proof of, a non-active site inhibition mechanism. The cooperative action of non-active site and catalytic site inhibitors increases the effectiveness of inhibition of an enzyme, and is a successful approach used with nucleoside and non-nucleoside inhibitors of HIV-1 reverse transcriptase (NRTI and NNRTI, respectively) in cART (33), as well as in cancer chemotherapy (34). Under the selective pressure of multiple PIs acting with different inhibitory mechanism, drug-naïve PRs would have more limitation to escape both classes of inhibitors. Although monotherapy regimes based on NNRTIs leads to resistance, the combination with NRTIs suppresses the evolution of RT resistance (11, 33). A similar scenario may be possible for PI-based therapy regimes if allosteric inhibitors would be delivered. There was no significant toxicity observed in PBMCs suggesting that with further chemical modification, these molecules may be delivered at pharmacological concentrations and tolerated by human cells. Overall, these non-active site PIs with multi-cell effectiveness represent promising approaches to expand the repertoire of current PIs to attenuate viral replication and prevent therapy resistance current PIs repertoire. Our study evaluated the impact of inhibitors in acute infection of primary lymphocytes, as the compounds were included on the day of infection. Future experiments are needed to test effect of non-active site PIs on longer term chronic infection in primary lymphocytes or latent cells.

## Acknowledgments

We want to thank Dr. Julie C. Williams for her help revising the manuscript. All authors declare no conflict of interest.

## Funding section

This study was supported in part by NIH R01 AI28571 (B.M.D., M.M.G. and S.P.); Stephany W. Holloway University Chair for HIV/AIDS Research (M.M.G.); and Laura McClamma Fellowship (C.M., R.M.C. and X.Z.).

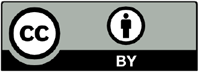 © 2019 by the authors. Submitted for possible open access publication under the terms and conditions of the Creative Commons Attribution (CC BY) license (http://creativecommons.org/licenses/by/4.0/).

## Mavian et al. Supplementary Information

**Fig S1.**
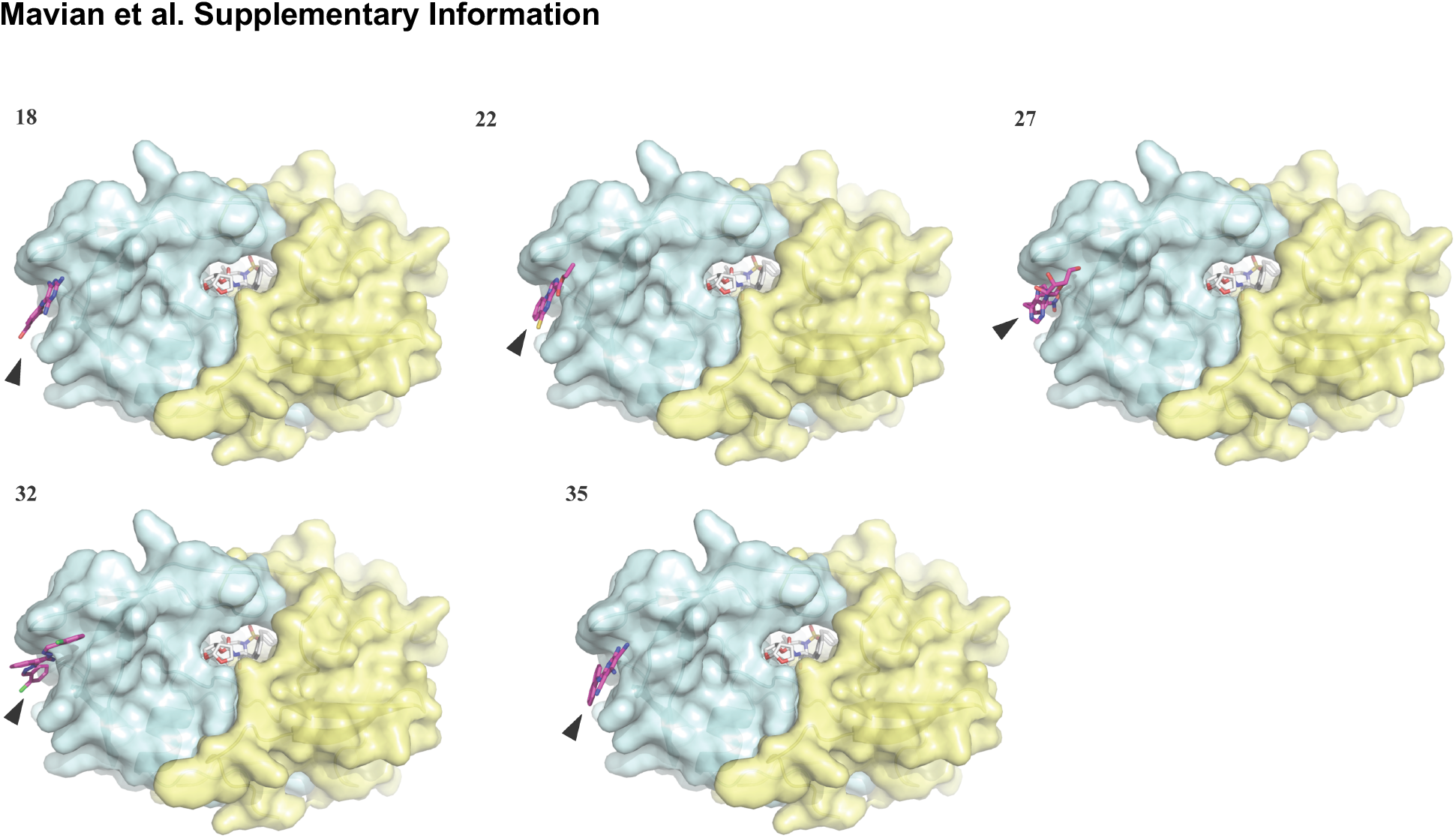
HIV PR subtype B wild-type in complex with compounds 18, 22, 27, 32 and 35. Computer modeling of compounds 18, 22, 27, 32 and 35 binding in the hinge region of the crystal structure of the PR homodimer complexed with inhibitor Darunavir in the active-site. Computer modeling of compounds, indicated by a black arrow, bound in the hinge region of the crystal structure of the PR homodimer complexed with PI inhibitor in the active-site (1t3r, http://www.rcsb.org/pdb/).

**Fig S2.**
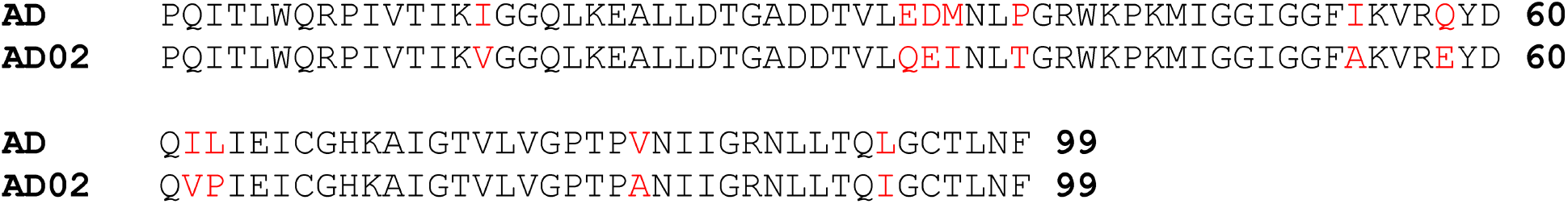
Amino acid alignment of the HIV-1_AD_ and HIV-1_AD02_ PRs. Amino acid alignment based on Clustal Omega (http://www.ebi.ac.uk/Tools/msa/clustalo/) of the wild-type and (AD) and drugresistant (AD02) PRs. Resistant residues that differ between HIV-1_AD_ and HIV-1_AD02_ are shown in red.

**Fig S3.**
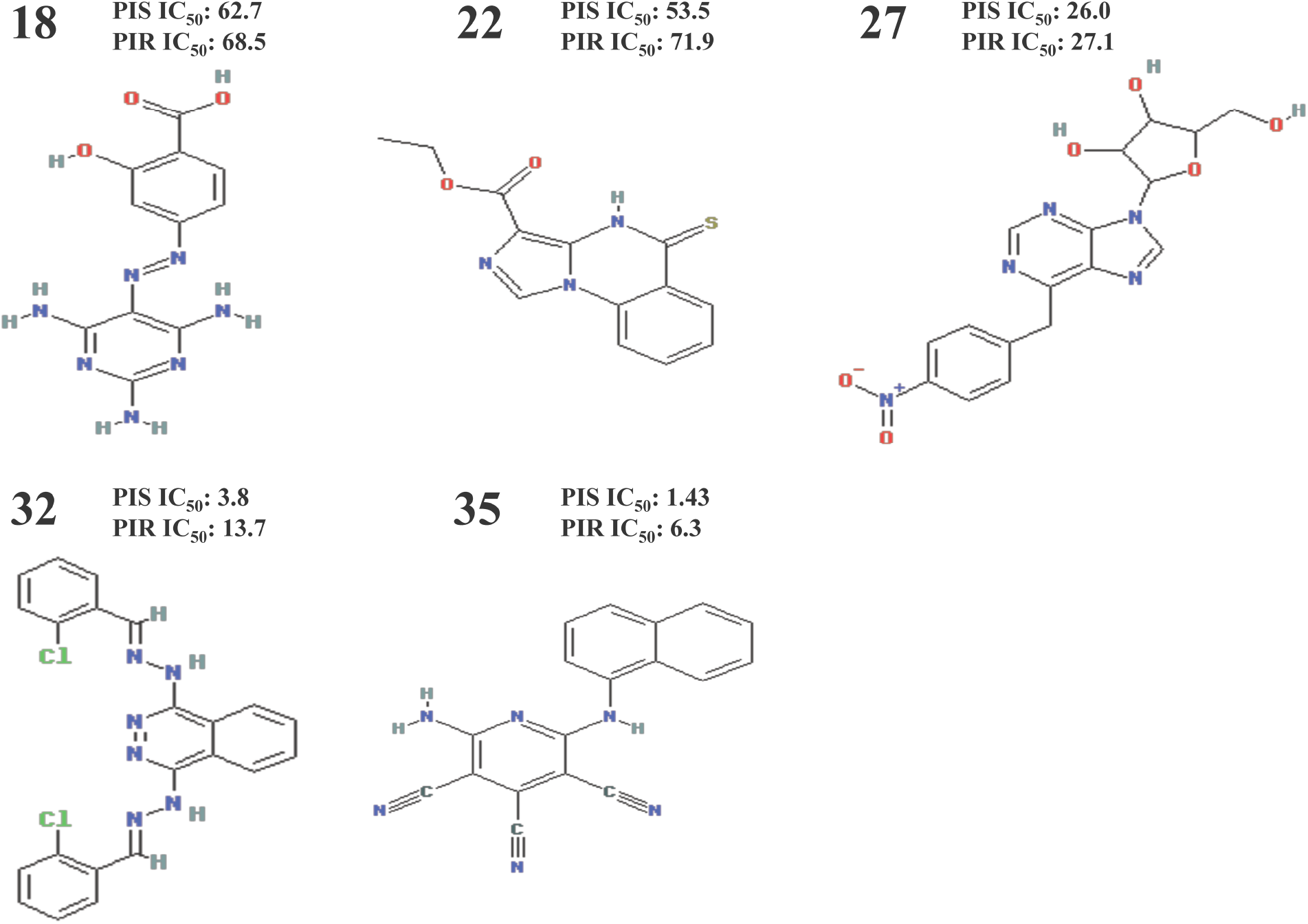
Chemical structure of the compounds. Chemical structure of the five lead compounds 18, 22, 27, 32 and 35. Next the identification number of each compound the IC_50_ against PI-sensitive [PIS IC_50_] or against multi PI-resistant HIV-1 variant [PIR IC_50_] are given in μM range.

**S1 Table.**
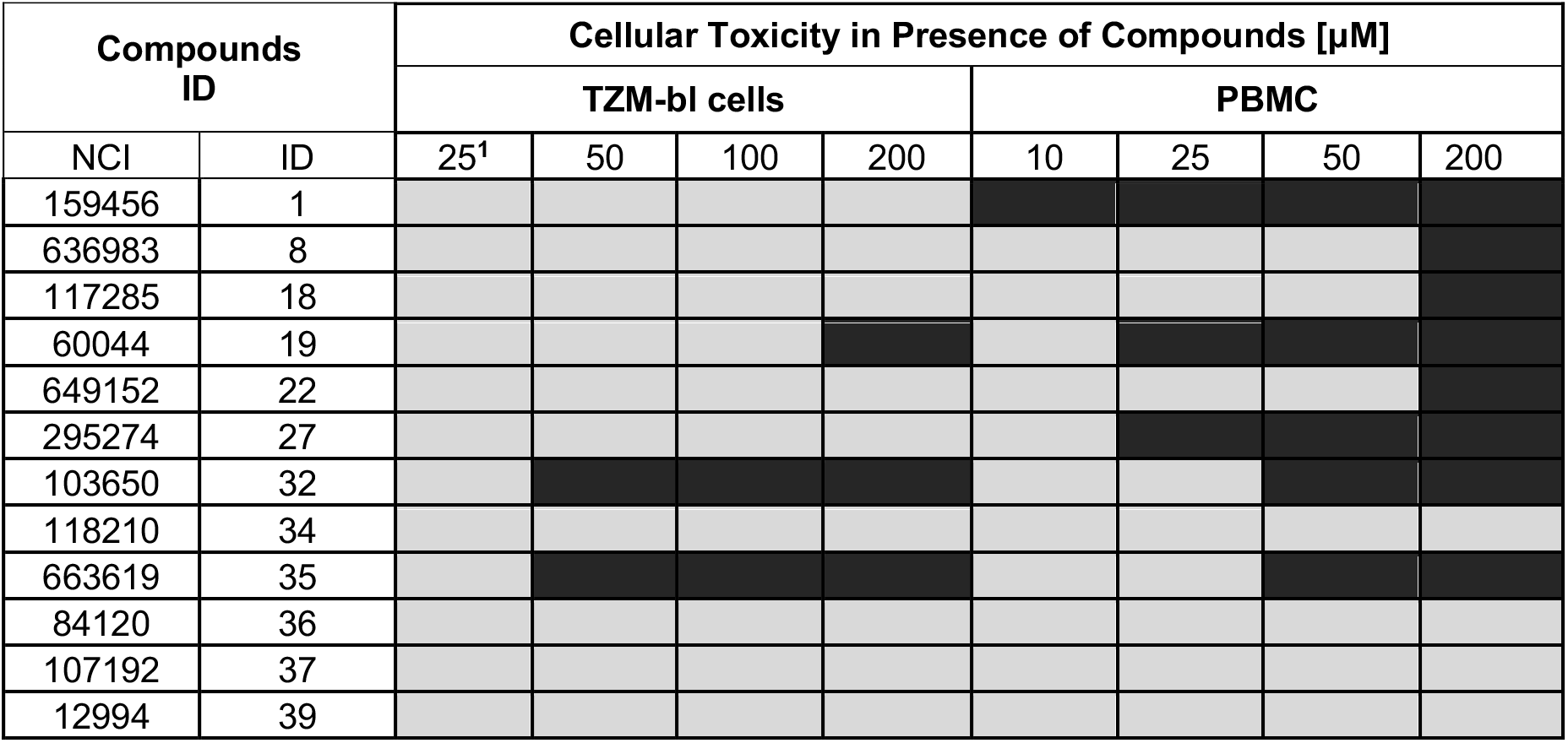
Toxicity of compounds in PBMC and TZM-bl cells. NCI: database number; ID: compound identification number; toxicity is expressed relative to viability of cells in DMSO: light grey, ≤10% or black, >10%.(^1^) Concentrations are expressed in μM.

**S2 Table.**
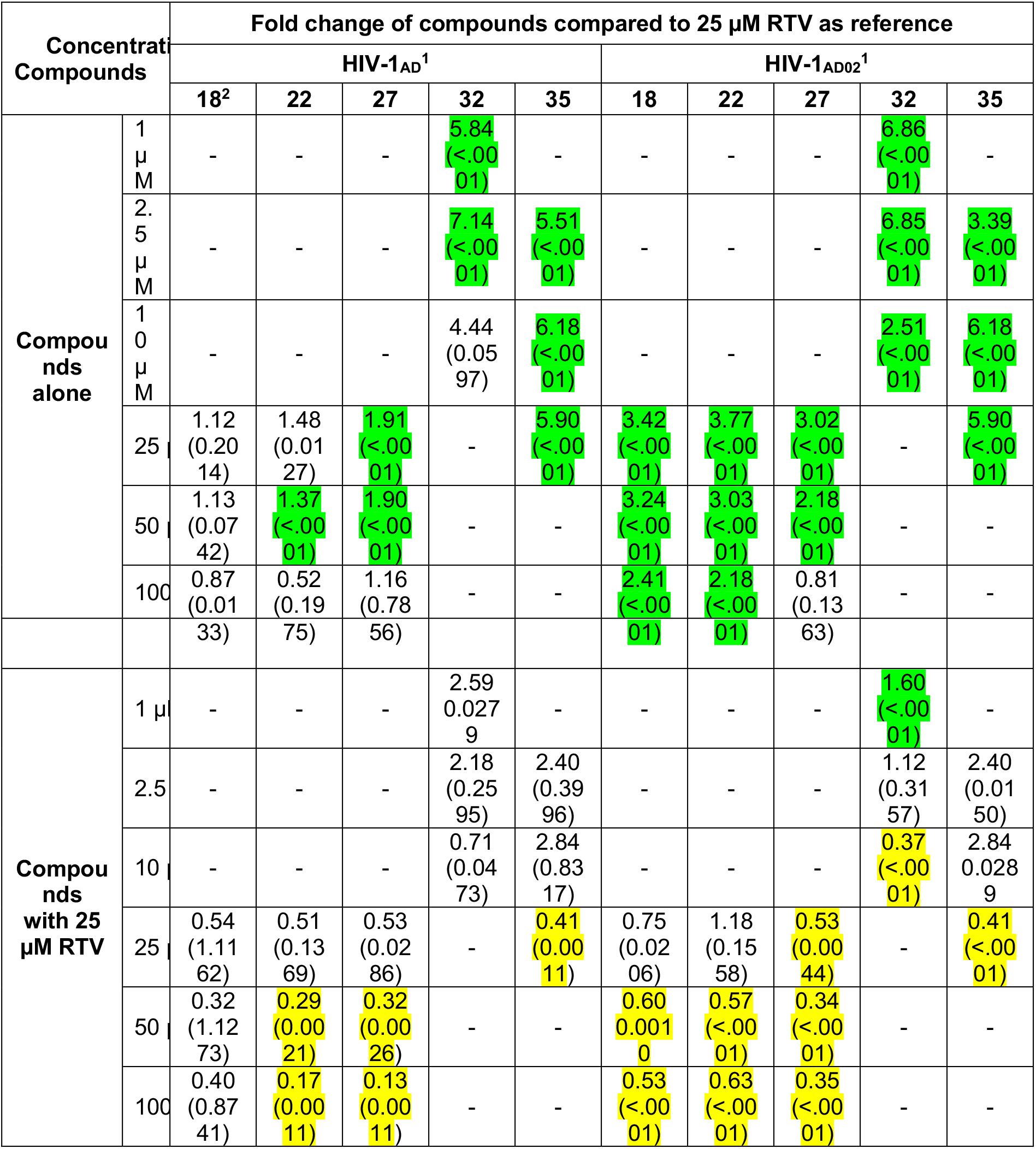
Fold change of compounds (18, 22, 27, 32 or 35) alone or in combination with 25 μM RTV compared to 25 μM RTV. P values are reported in parenthesis with Bonferroni correction with significance p<0.0055; fold change >1 indicates antiviral-activity less effective (viral replication higher) than 25 μM RTV alone and significant ones are highlighted in green; fold change <1 indicates antiviral-activity more effective (viral replication lower) than 25 μM RTV alone and significant ones are highlighted in yellow; (-) not performed. (^1^) Anti-viral activity against HIV-1; (^2^) Compounds identification number.

## Notes

### Competing Interest Statement

The authors have declared no competing interest.

